# Locus coeruleus activity correlates with transient elevations in breathing during opioid-induced respiratory depression

**DOI:** 10.64898/2026.11.02.763547

**Authors:** Caroline C. Szujewski, Jaseph Soto-Perez, Anmin Yang, Matthew Warden, Shivang Sullere, Gia E. Fisher, Joseph V. Gogola, Alfredo J. Garcia

## Abstract

A history of use remains a key risk factor for opioid-related overdose deaths in the ongoing opioid epidemic. The locus coeruleus (LC) is the principal source of noradrenergic tone and a site for opioid neuromodulation and tolerance. How a history of opioid use affects LC activity and shapes opioid-induced respiratory depression (OIRD) remains poorly understood. Using plethysmography and fiber photometry, we assessed how fentanyl affects LC activity and OIRD before and after five days of repeat opioid use (ROU). Fentanyl increased the frequency of LC bursts independent of exposure history and was coupled to transient elevations in breathing. LC coupled ventilation (LC-Ve) persisted through ROU. After ROU, OIRD magnitude diverged across mice. Mice developing an adaptive ventilatory response (AVR) showed a smaller OIRD that coincided with improved ventilation outside LC bursts and reduced LC-Ve gain. Mice with the non-adaptive response (NVR) show no improvement in OIRD or ventilation outside LC bursts, resulting in an unchanged reliance on LC-Ve. While LC-Ve reflexively counters OIRD acutely, it may also drive oscillations to destabilize blood gas homeostasis, perpetuating pathophysiology with repeat use. Thus, LC-coupled breathing appears to be a physiological signature of divergent respiratory responses with a history of fentanyl exposure.

**Impact statement:** Noradrenergic neurons of the locus coeruleus are activated by fentanyl and coincide with brief surges in breathing during opioid-induced respiratory depression. Reliance on these surges distinguishes mice that fail to develop ventilatory tolerance.

## Introduction

The United States continues to grapple with an opioid epidemic that has claimed over 800,000 lives since 1999 (CDC, 2023). Opioid overdose begins with opioid-induced respiratory depression (OIRD), the first step in the pathophysiological cascade leading to death. Substantial progress has been made toward understanding how opioids act on discrete brainstem respiratory centers to produce OIRD (Bachmutsky et al., 2020; Varga et al., 2020; Baldo and Rose, 2022), but these advances have largely addressed regional pharmacological actions rather than network-level phenomena. A growing body of evidence in rodents (Laferrière et al. 2005, Hou et al. 2023, Rai et al., 2025) and humans (Algera et al. 2021) indicates that persistent opioid exposure reshapes ventilation. This is prominently evidenced by the emergence of an adaptive ventilatory response (AVR) where the magnitude of OIRD is diminished (Laferrière et al. 2005, Rai et al., 2025) and improved ventilatory-metabolic coupling during opioid challenge (Rai et al., 2025). While these observations indicate that repeat opioid use (ROU) produces OIRD tolerance, the phenomenon does not develop uniformly across individuals (Rai et al. 2025) and can be augmented by contextual cues that promote associative learning (Hou et al. 2023). These phenomena may contribute to the observed variability in OIRD susceptibility among individuals and across occasions of use (Monforte, 1977; Fox et al., 2018; Bubier et al., 2020; Dong et al., 2020). As opioid signaling is known to affect neuromodulatory systems throughout the brain (Aghajanian 1978, Rasmussen et al. 1990, Williams et al. 2001), ROU-dependent changes in neuromodulatory activity may determine who develops tolerance and how durably.

The locus coeruleus (LC), the principal source of noradrenergic neuromodulation in the brain, is well positioned to act as a neural substrate shaping adaptations in response to ROU. LC neurons regulate breathing and both project to and receive input from higher-order brain areas and the brainstem (Aston-Jones et al., 1986; Ennis et al., 1998; Samuels and Szabadi, 2008), including cardiorespiratory control centers (Westlund and Coulter, 1980, Ter Horst et al. 1991, Dobbins and Feldman, 1994, Biancardi et al. 2008). Functionally, the LC has been implicated in vigilance, arousal, stress responses, attention to salient stimuli, memory, and autonomic regulation (Elam et al., 1984; Aston-Jones et al., 1991; Aston-Jones and Cohen, 2005; Valentino and Van Bockstaele, 2008; Carter et al., 2010; Hansen, 2017). Critically, its output is shaped by prior experience, with LC neurons remodeling their firing as reward contingencies change (Bouret and Sara, 2004) and being required for acquisition of conditioned cardiovascular responses (Harris and Fitzgerald, 1991). The LC therefore sits at a neurophysiological intersection between experience-dependent plasticity and the brainstem networks that generate breathing.

The LC is also a canonical site of opioid action. All three opioid receptor classes are present in the LC (van Bockstaele et al. 1997; Ding et al. 1996, Al-Hasani et al. 2013), and acute opioid exposure hyperpolarizes LC neurons and suppresses their firing (Pepper and Henderson, 1980; Christie, 1991). With chronic exposure, LC neurons become tolerant to morphine and display withdrawal hyperexcitability (Aghajanian 1978). However, these adaptations have been characterized largely in slice and anesthetized preparations and in relation to dependence and withdrawal, leaving unresolved what the LC does in awake animals during OIRD.

Here we test how ROU shapes in vivo LC activity and its relationship to breathing during fentanyl challenge in naive and opioid-experienced mice. Furthermore, we assess whether such relationships depend on the context of administration. Using simultaneous whole-body plethysmography and fiber photometry, we monitored breathing and LC activity in awake behaving mice across five days of ROU, with fentanyl being delivered in a drug-paired (FP) environment and saline in a drug-unpaired (SP) environment. We find that fentanyl stimulates rather than silences LC activity in vivo, and that this activity is coupled to transient elevations in ventilation (LC-Ve) that periodically stimulate breathing during OIRD on both Day 1 and Day 5 of ROU. ROU produced an adaptive ventilatory response (AVR), characterized by improved ventilation outside LC events and reduced LC-Ve gain, and a non-adaptive response (NVR), which lacked improved breathing outside LC activity and continued to rely on LC-Ve. While the context of administration did not change the magnitude of depression, it shaped the pattern of OIRD. These findings indicate that LC activity is coordinated with reflexive attempts to stimulate breathing during OIRD and contributes to heterogeneous breathing patterns in response to fentanyl.

## Results

To explore how repeat opioid use (ROU) influences LC activity and OIRD, breathing and LC GCaMP fluorescence were simultaneously measured in awake-behaving adult mice before and after fentanyl administration for 5 days of ROU (Figure 1Ai-iii). When mice were opioid-naive (i.e., Day 1 of ROU), we observed transient bursts of LC activity that coincided with periods of increased breathing (LC-Ve). Breathing during LC-Ve exhibited increased minute ventilation (Figure 1C, paired t test, p = 0.026), respiratory rate (Figure 1D, paired t test, p = 0.001), and tidal volume (Figure 1E, paired t test, p = 0.003). LC-Ve occurred both before and during fentanyl administration. While fentanyl caused a persistent ventilatory depression (Figure 1F), LC burst frequency increased (Figure 1G, paired t test, p = 0.002) as did LC-Ve events, which reflexively stimulate breathing during OIRD despite their transient nature (Figure 1H, paired t test, p = 0.002).

**Figure 1.**
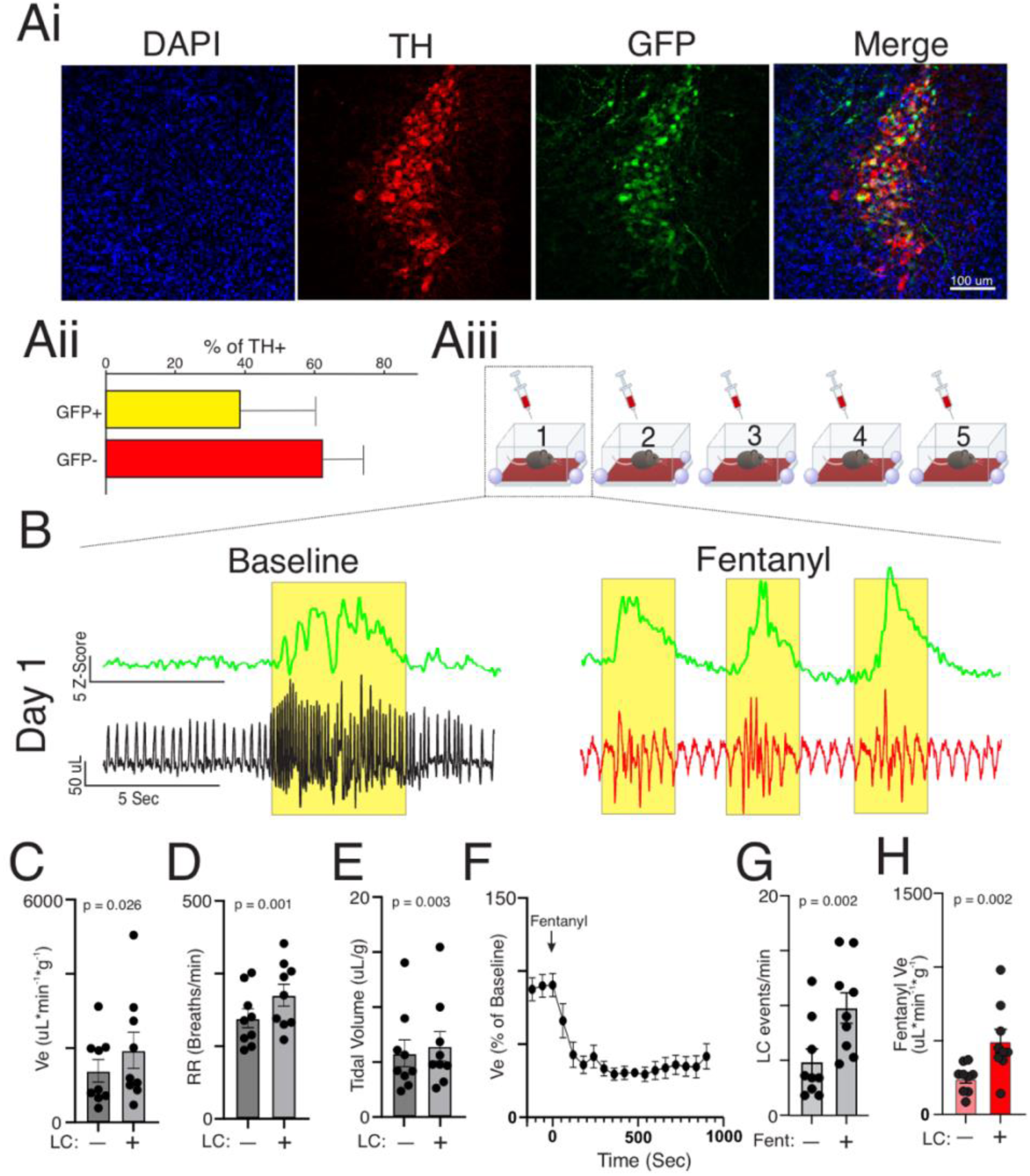
LC activity correlates with transient elevations in ventilation during baseline and OIRD. (Ai) Representative immunohistochemistry of the LC from a mouse following AAV-GCaMP8 expression shows expression of GFP in TH+ cells. (Aii) Quantification of GCaMP expression within TH+ cells of the LC from experimental mice (n = 9). (Aiii) Experimental schematic showing recording of locus coeruleus activity and whole-body plethysmography measurements of breathing in awake mice throughout ROU protocol. Breathing and LC activity were recorded prior to fentanyl administration (i.e., baseline conditions) and after fentanyl administration. (B) Representative traces of LC activity (green) and corresponding breathing traces during baseline breathing (black) and OIRD (red) on Day 1 of ROU (i.e., when mice are naive to fentanyl). Shaded regions (yellow) indicate LC events. During baseline, LC events coincide with transient increases in respiratory activity. Quantification of baseline (C) Ve, (D) respiratory rate and (E) tidal volume with (LC+) and without (LC−) detected LC bursting (n = 9; C, p = 0.026; D, p = 0.001; E, p = 0.003). LC+ coincides with transient increases in breathing under baseline conditions before fentanyl administration. After fentanyl administration (red), periodic LC activity remains active and linked to transient increases in breathing during OIRD. (F) Time course of fentanyl-induced respiratory depression, illustrating the rapid and sustained nature of depressed ventilation caused by the opioid. (G) The number of LC burst events is increased during OIRD (p = 0.002). (H) Quantification of Ve during OIRD with (LC+) and without (LC−) LC bursting (p = 0.002). During LC+, Ve is larger than during LC−, indicating that coupling between transient LC activation and ventilation stimulation persists even during fentanyl challenge. (C-E, G, H) Paired t-test. All summary data presented as mean with SEM.

Across all subjects tested, the ventilatory response (Ve) to fentanyl challenge differed between Day 1 and Day 5 of ROU (Figure 2A, Mixed Effects analysis, D1 vs D5: p = 0.023), yet LC activity under fentanyl was not significantly different following ROU (LC events per minute, D1 vs D5; p = 0.13; paired t-test, n = 9). A greater mean Ve was maintained under fentanyl on Day 5 compared to Day 1 (Figure 2Bi, paired t test, p = 0.023), consistent with the previously reported improvement of breathing during OIRD following ROU (Rai et al., 2025). As OIRD is known to be variable even following ROU (Rai et al. 2025), we next calculated the OIRD Ratio Index (ORI) for all individual to demonstrate that improved ventilation during OIRD was not uniform acrross the mice. Most mice (n = 6 of 9) an exhibited an adaptive ventilatory response (AVR, ORI > 0). Whereas a minority (n = 3) exhibited the non-adaptive ventilatory response (NVR, ORI ≤ 0) (Figure 2Bii, unpaired t test, p = 0.017). Upon endotype segregation, AVR mice showed improved breathing during OIRD on Day 5 (Figure 2Ci-iii), while NVR mice showed no change in OIRD after ROU (Figure 2Di-iii). Together, these data indicate that ROU can promote ventilatory tolerance to fentanyl-induced respiratory depression, but this response is heterogeneous across subjects.

**Figure 2.**
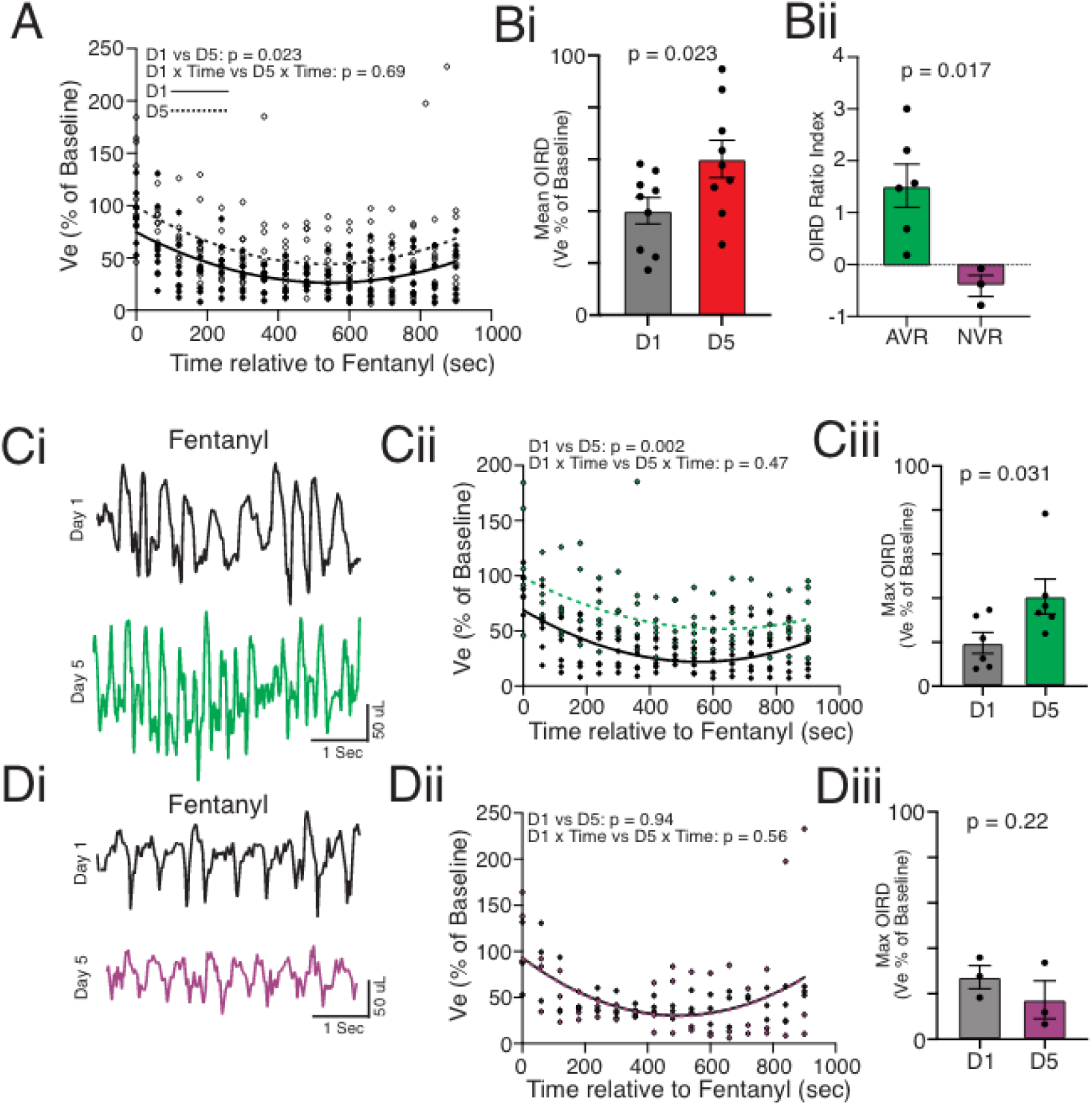
Repeat opioid use reveals heterogeneous ventilatory adaptations during fentanyl-induced respiratory depression. (A) Time course of the Ve response to fentanyl on Day 1 (D1: solid line; black circles represent individual responses) and Day 5 (D5: dotted line; open circles represent individual responses) of ROU showing differences between days (D1 vs D5, p = 0.023, n = 9). (Bi) Comparing the average depression of Ve during OIRD when mice were experienced with ROU (i.e., Day 5) to when they were naive (i.e., Day 1) revealed that experience with ROU leads to a greater Ve during OIRD, consistent with the development of an adaptive ventilatory response to OIRD (p = 0.023, n = 9). (Bii) Consistent with observations previously reported by Rai et al. (2025), examination of the individual ORI revealed that the majority of individual response profiles (n = 6 of 9) showed the AVR. However, a minority of mice (n = 3 of 9) exhibited a non-adaptive ventilatory response to ROU (NVR) (p = 0.017). (Ci) Representative breathing trace from an AVR mouse showing improved breathing during fentanyl challenge by Day 5 of ROU. (Cii) Time course of Ve response to fentanyl on D1 (black) and D5 (green) from AVR mice demonstrated greater Ve during OIRD after ROU (D1 vs D5, p = 0.002, n = 6). (Ciii) Summary data of minimum Ve during OIRD in AVR mice on Day 1 (black) and Day 5 (green) (n = 6, p = 0.031). (Di) Representative breathing trace from an NVR mouse showing no improvement in breathing by Day 5 of ROU. (Dii) Time course of Ve response to fentanyl on D1 (black) and D5 (purple) from NVR mice demonstrating no improvement of Ve during OIRD after ROU (D1 vs D5, p = 0.94, n = 3). (Diii) Summary data of minimum Ve during OIRD in NVR mice on Day 1 (black) and Day 5 (purple) (n = 3, p = 0.22). (Bi, Diii) Paired t-test. (Bii) unpaired t test. (Ciii) Wilcoxon test. (A) Mixed Effects with Tukey’s multiple comparison. (Cii, Dii) RM two-way ANOVA with Tukey’s multiple comparisons and Geisser-Greenhouse correction. All summary data presented as mean with SEM.

Adaptive changes in sympathetic signaling have been reported following ROU in both murine models and humans (Kienbaum et al., 2001; Akbarian et al., 2002). While these studies establish the possibility that brain norepinephrine signaling may change with ROU and potentially contribute to OIRD tolerance, this has yet to be tested. Given its role as the primary source of adrenergic signaling in the brain, we tested this hypothesis by measuring LC activity during OIRD prior to and following ROU. LC activity following ROU was consistent with that observed when mice were naive (Figure 3A), as LC-Ve events were present in baseline (Figure 3B, Wilcoxon test, p = 0.004) and during OIRD (Figure 3C, paired t test, p = 0.019), and frequency of LC events increased during OIRD (Figure 3D, paired t test, p = 0.035). In AVR mice, max OIRD (minimum Ve during fentanyl exposure normalized to baseline) outside of LC activity was improved on Day 5 of ROU as compared to Day 1 (Figure 3E, Two-way ANOVA with Fisher’s LSD, p = 0.009). In contrast, LC-Ve was reduced following ROU (Figure 3G, p = 0.001). In NVR mice, max OIRD outside of LC activity was similar on Day 1 and Day 5 of ROU, suggesting a lack of OIRD tolerance (Figure 3E, Two-way ANOVA with Fisher’s LSD, p = 0.54). Additionally, there was no change in magnitude of LC-Ve following ROU (Figure 3H). While LC-Ve was not different between respiratory endotypes when the groups were initially exposed to fentanyl on Day 1, NVR mice displayed elevated LC-Ve activity compared to AVR mice following ROU (Supplement Figure 1, p = 0.035). Emergence of different tolerance modalities between AVR and NVR groups was further evidenced by a significant difference in the Day-Group interaction (Figure 3I, p = 0.002). These results suggest that tolerance to OIRD is driven by elevated Ve outside of LC activity, and therefore the LC-associated Ve becomes less effective against OIRD following ROU. When this non-LC-mediated form of tolerance fails to develop, as observed in NVR mice, reliance on reflexive LC-driven Ve remains the principal form of tolerance to OIRD.

**Figure 3.**
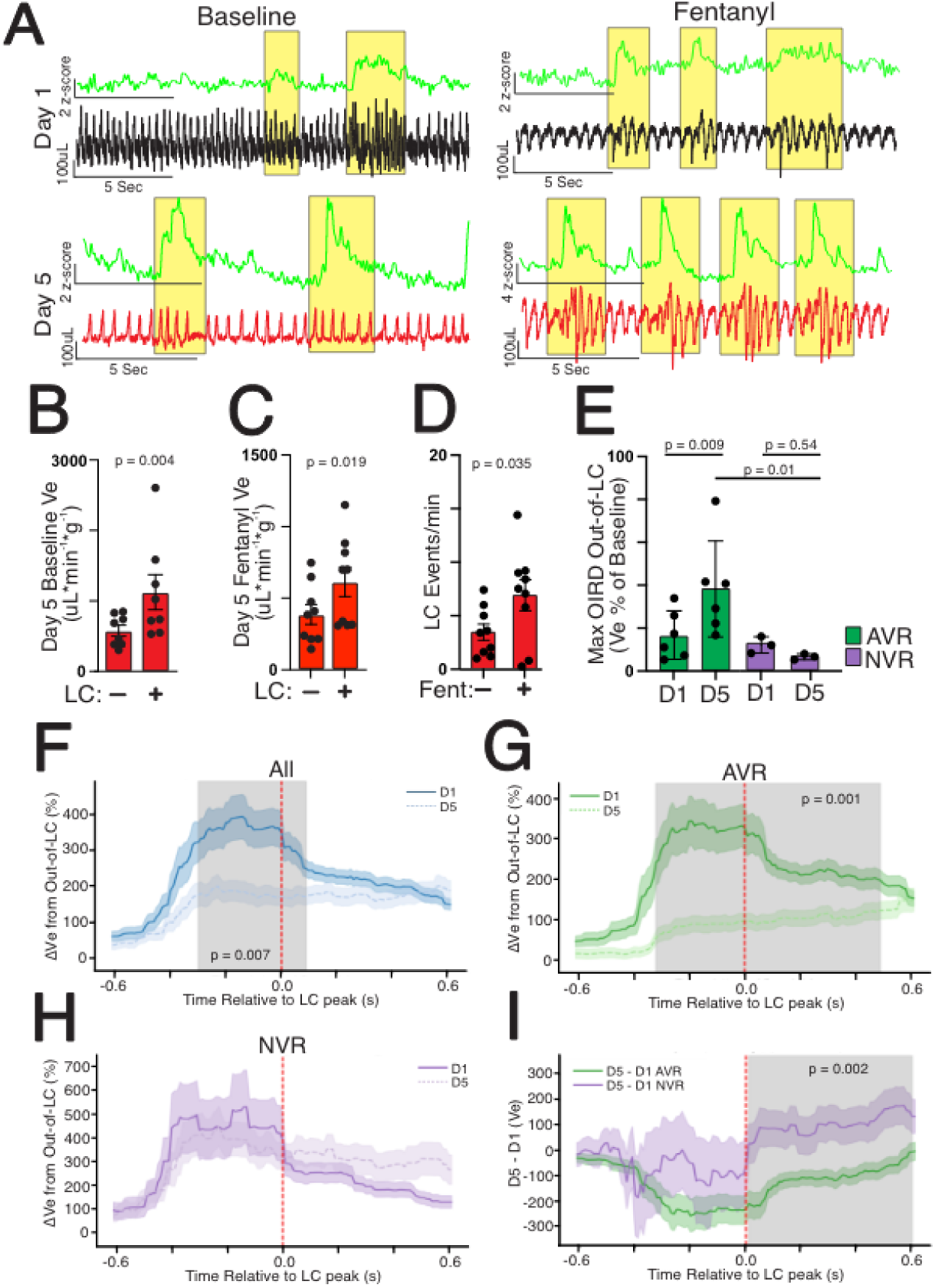
Repeated opioid use does not abolish locus coeruleus recruitment during fentanyl-induced respiratory depression. (A) Representative traces of Z-scored LC GCaMP activity (green) and breathing on Day 1 (black) and Day 5 (red) of ROU, showing that LC activity during baseline conditions and in fentanyl remains qualitatively similar to responses observed in opioid-naive mice, where bursts of transient LC activity (highlighted in yellow) coincide with increased breathing. (B) Summary data of Ve in and outside of LC GCaMP events shows transient increases in Ve occur on Day 5 of ROU in baseline (n = 9, p = 0.004). (C) Summary data of Ve in and outside of LC GCaMP events shows transient increases in Ve occur on Day 5 of ROU under fentanyl (n = 9, p = 0.019). (D) On Day 5 of ROU, frequency of LC GCaMP bursts was increased under fentanyl (n = 9, p = 0.035). (E) Summary data of maximum OIRD (minimum Ve during fentanyl exposure normalized to baseline) outside LC events shows Ve outside LC events was greater on Day 5 of repeated opioid use in AVR mice (n = 6, p = 0.009). On the other hand, summary data from NVR mice show that max OIRD Ve outside of LC activity does not change (p = 0.54) following ROU (n = 3). (F) Time course of Ve relative to peak LC activity on D1 (solid line) and D5 (dotted line) from all mice, showing a decrease in the LC-Ve relationship under fentanyl following ROU (p = 0.007, n = 9). (G) Time course of Ve relative to peak LC activity on D1 (solid line) and D5 (dotted line) from AVR mice (n = 6) during OIRD (p = 0.001). (H) Time course of Ve relative to peak LC activity on D1 (solid line) and D5 (dotted line) from NVR mice (n = 3) during OIRD. (I) Interaction between AVR and NVR groups across Day 1 and Day 5, demonstrating that the groups showed different response patterns (p = 0.002, AVR n = 6, NVR n = 3). Shaded area indicates periods of significant (p < 0.05) difference between D1 and D5. (B) Wilcoxon test. (C-D) Paired t-test. (E) Two-way ANOVA with Fisher’s LSD. (F-I) Linear mixed-effects model, significance determined by cluster-based permutation test. All summary data presented as mean with SEM.

Based on observations supporting the concept of contextual differences in overdose-related mortality (Siegel, 1978; Siegel et al., 1982), we next tested whether a previous association of an environmental context with fentanyl affected respiratory and LC activity. We tested respiratory and LC activity in response to fentanyl in a previously fentanyl-paired environment (FP) and a previously saline-paired environment (SP). The pattern of respiratory depression was found to differ when fentanyl was administered in the SP environment (Figure 4Aii, RM two-way ANOVA with Tukey’s multiple comparisons, SP x Time vs FP x Time: p = 0.014). However, neither Fentanyl Ve outside of LC activity (Figure 4Aiii, p = 0.91) nor mean ventilatory depression (Figure 4Aiv, paired t test, p = 0.67) differed across context. During OIRD, LC activity was also similar in frequency (LC events per minute, SP vs FP; p = 0.83; paired t-test, n = 9) and LC-Ve transiently increased in both contexts (Figure 4Bii-iii). Upon stratifying the mice into endotypes (Figure 5A), LC-Ve events were transiently stimulated in both AVR (Figure 5Bi) and NVR (Figure 5Bii) mice independent of the context of drug delivery. In AVR mice, LC-Ve was similar in both SP and FP contexts (Figure 5Bi), whereas in NVR mice, Ve remained consistently elevated following the peak of the LC burst (Figure 5Bii, p = 0.03). Despite LC-Ve differing between SP and FP during a post-peak interval in NVR mice, the corresponding context-by-response-group interaction effect for this period did not survive cluster-level correction (Figure 5Biii; 0.03 - 0.27 s, p = 0.07). Furthermore, we found no difference in LC-Ve between AVR and NVR groups in either the SP or FP contexts (Supplement Figure 2). Thus, while the simple-effect analysis indicates context-dependent modulation in NVR mice (Figure 5Bii), the interaction analysis provides only suggestive evidence that this modulation differs by response group. No differences were found in max OIRD Ve outside of LC across contexts in either AVR or NVR mice (Figure 5C, RM two-way ANOVA with Fisher’s LSD, Context factor: p = 0.9).

**Figure 4.**
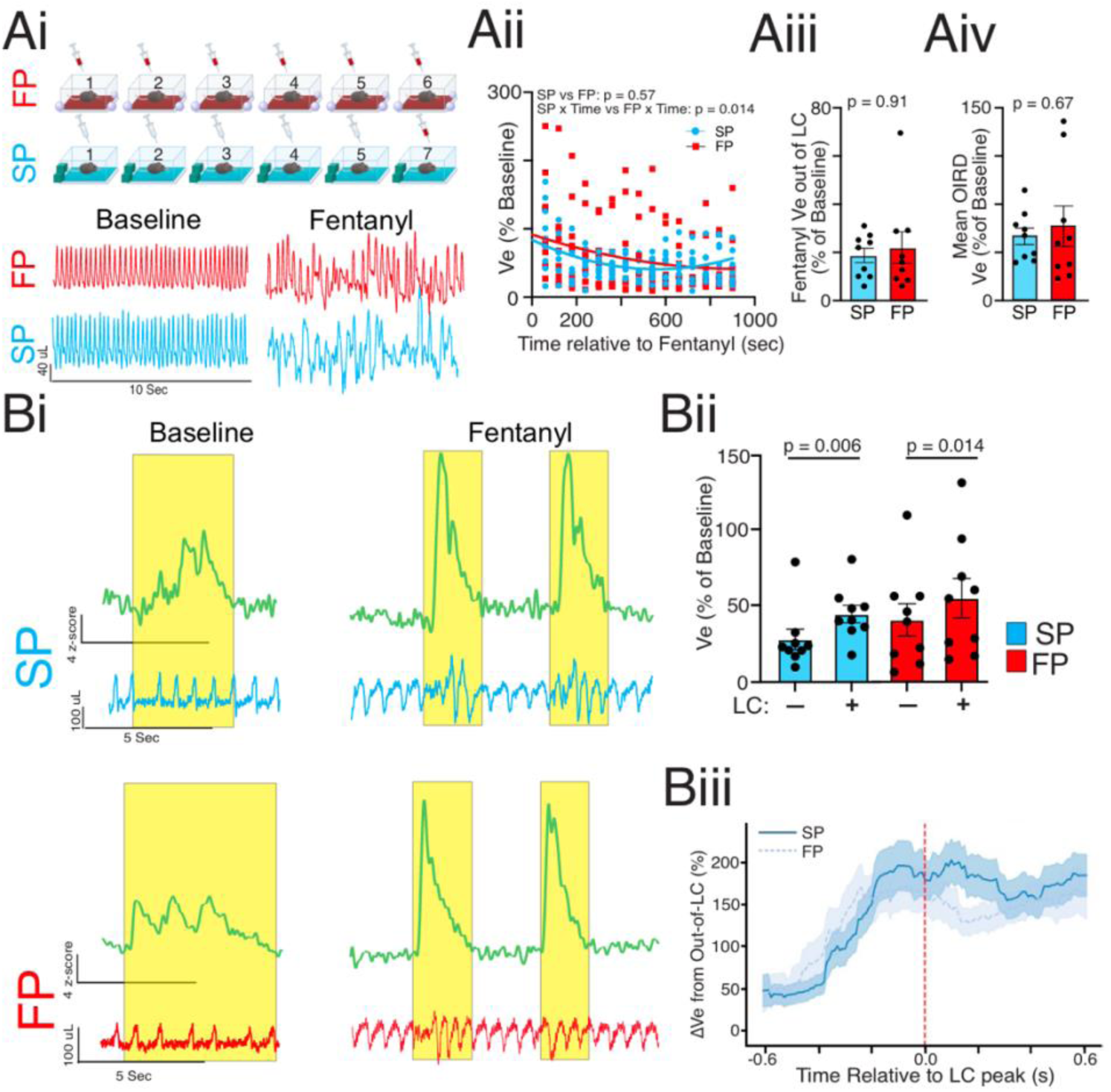
Context does not shape the severity of OIRD following ROU. (Ai) Top: schematic of the 5-day experimental ROU protocol shows daily injection of fentanyl in the drug-paired context (FP, red) and saline in the drug-unpaired context (SP, cyan) for five consecutive days. On days 6 and 7, mice received fentanyl in either context (FP or SP) one day and fentanyl again on the following day in the opposite context. Bottom: representative respiratory traces under baseline (left) and fentanyl (right) in the fentanyl-paired (FP, top) and saline-paired (SP, bottom) environments. (Aii) Time course of Ve response to fentanyl (normalized to baseline) in the SP (cyan) and FP (red) environments shows OIRD pattern differs across contexts (n = 9; SP × Time vs FP × Time, p = 0.014). (Aiii) Summary data of mean OIRD Ve outside of LC events in the SP (cyan) and FP (red) environments (n = 9, p = 0.91). (Aiv) Comparing mean OIRD between SP and FP environments shows no difference in Ve depression across contexts (n = 9, p = 0.67). (Bi) Representative traces of Z-scored GCaMP activity (green) and ventilation in the SP (cyan) and FP (red) conditions. (Bii) Summary data of ventilation in (+) and outside (-) of LC GCaMP activity shows Ve is stimulated under LC activity in both SP (p = 0.006) and FP (p = 0.014) environments (n = 9). (Biii) Time course of Ve relative to peak LC activity in the SP (solid line) and FP (dotted line) from all mice (n = 9) during OIRD. (Aii, Bii) RM two-way ANOVA with Tukey’s multiple comparisons. (Aiii) Wilcoxon test. (Aiv) Paired t-test. (Biii) Linear mixed-effects model, significance determined by cluster-based permutation test. All summary data presented as mean with SEM.

**Figure 5.**
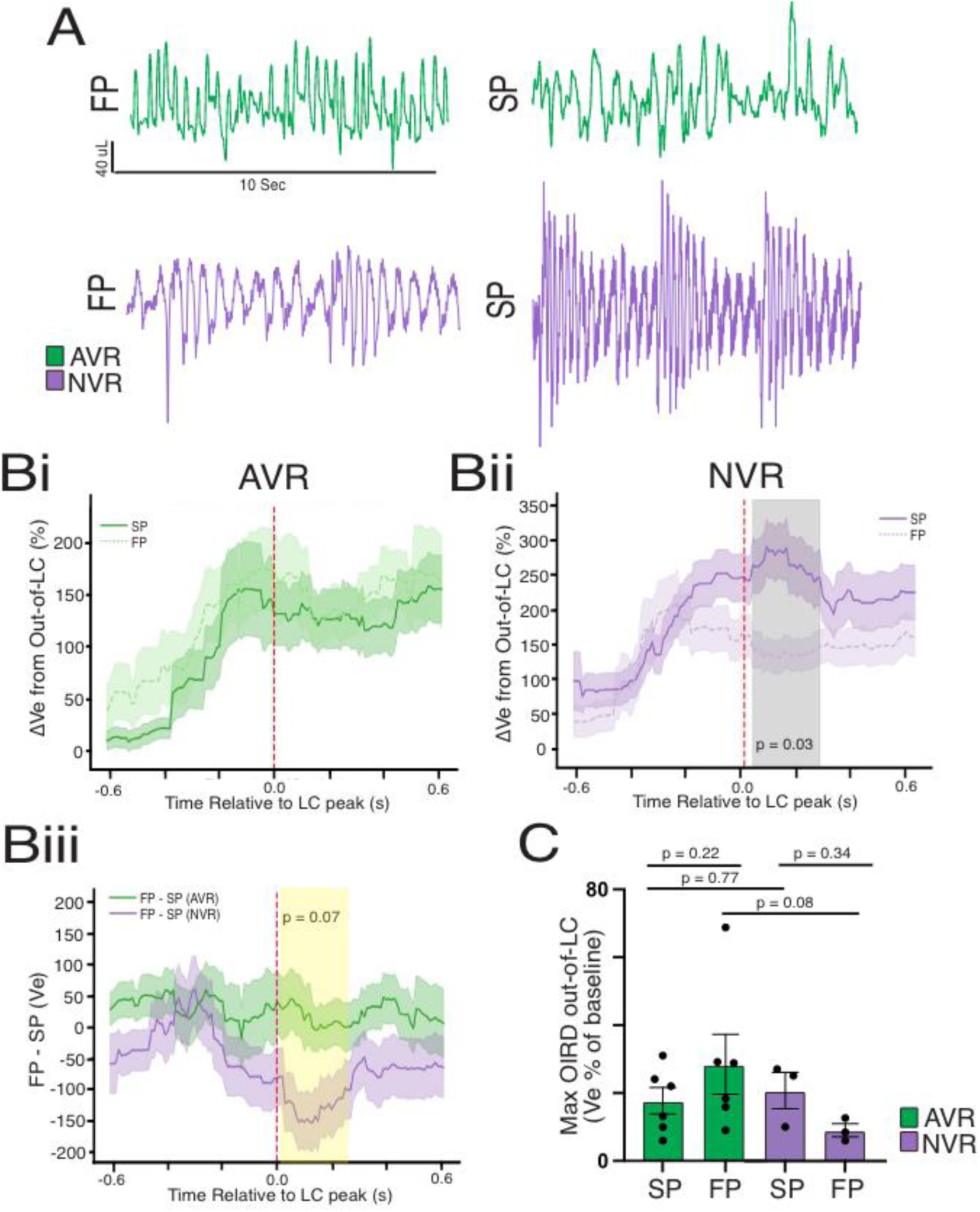
Evaluating divergent contextual tolerance mechanisms in OIRD endotypes. (A) Representative respiratory traces from AVR (top) and NVR (bottom) mice under fentanyl in the FP (left) and SP (right) environments. (Bi) Time course of Ve relative to peak LC activity in the SP (solid line) and FP (dotted line) from AVR mice (n = 6) during OIRD. (Bii) Time course of Ve relative to peak LC activity in the SP (solid line) and FP (dotted line) from NVR mice (n = 3) during OIRD (p = 0.03). (Biii) Interaction between AVR and NVR groups across SP and FP contexts during OIRD (0.03-0.27 s, p = 0.07). (C) Summary data of maximum OIRD (minimum Ve during fentanyl exposure normalized to baseline) in the SP and FP environments from AVR (n = 6) and NVR (n = 3) mice. Gray shaded areas indicates significant difference between groups. Yellow shaded areas indicates where significant difference between groups was tested but did not reach significance. (Bi-iii) Linear mixed-effects model, significance determined by cluster-based permutation test. (C) RM two-way ANOVA with Fisher’s LSD. All summary data presented as mean with SEM.

## Discussion

Respiratory depression is the hallmark of opioid overdose and the principal driver of the pathophysiological cascade of events that, when left untreated, can end in death. While there is increasing recognition that persistent opioid use remodels respiratory control, the neurophysiological changes occurring with ROU remain poorly resolved. Using a preclinical model of repeat opioid use (ROU), we characterized how repeated fentanyl exposure and the context of drug administration affect ventilation and LC activity during OIRD. We find that fentanyl causes transient elevations in ventilation that are often coupled to LC activity (LC-Ve). Together, our findings frame LC-coupled respiratory patterns as reflexive compensatory responses to OIRD, whose strength and attributes vary across AVR and NVR endotypes and may be sensitive to the context of drug administration.

The LC is the principal source of brain noradrenaline and supports central nervous functions such as respiratory control (Krohn et al., 2023), arousal (Ross and Van Bockstaele, 2021), and salience detection (Sara and Bouret, 2012). It is uniquely positioned as a central hub sending and receiving input to and from relevant brain areas that may influence the opioid overdose trajectory. Local acidification of the LC stimulates breathing, while lesioning the LC depresses the hypercapnic ventilatory response (de Carvalho et al., 2014; Biancardi et al. 2008; Gargaglioni et al., 2010). This may involve recruitment of downstream networks that contribute to the drive to breathe, such as the Kölliker-Fuse nucleus, which receives both glutamatergic and noradrenergic drive from the LC (Varga et al., 2026). The LC also exerts influence over cortical activation and sleep-to-wake transitions (Carter et al., 2010), and receives direct input from preBötzinger neurons to promote arousal (Yackle et al., 2017). Furthermore, LC activation facilitates respiratory rhythm generation through noradrenergic signaling (Hilaire et al., 2004). Thus, the LC may act as a critical node in OIRD as it coordinates noradrenergic-arousal signaling throughout the respiratory circuit.

Because the LC expresses mu opioid receptors, fentanyl should suppress its intrinsic excitability (Pepper and Henderson, 1980; Christie, 1991). However, our experiments showed that LC activity is stimulated by fentanyl in vivo, and this activity temporally coincides with transient elevations in breathing during OIRD. Whether LC activity is upstream, downstream, or in parallel with respiratory circuitry driving LC-Ve remains unknown. However, optogenetic activation of Tac1 neurons in the parabrachial nucleus (PB) drives high-frequency breathing (Arthurs et al., 2023) reminiscent of LC-Ve. Given that the LC sends noradrenergic and glutamatergic projections to the PB (Yang et al., 2021), connectivity between the LC and the PB may be involved in the generation of LC-Ve during OIRD.

Acutely, LC-Ve may reflexively counter OIRD and the progression toward terminal apnea. However, the persistent transitory pattern of LC-Ve also causes waxing and waning of ventilation similar to oscillatory breathing patterns as found in Cheyne-Stokes breathing (Leung et al., 2003) and Biot’s breathing (Webber and Speck, 1981) which drive instability in blood gas homeostasis and can lead to sympathetic surges that can be independently pathogenic (van de Borne et al., 1998; Lorenzi-Filho et al., 1999; Brack et al., 2012; Leuenberger et al., 1995). Given evidence of LC chemosensory capacity, blood-gas oscillations may be contributing to LC bursting (Elam et al. 1981), which may consequently stimulate breathing during OIRD. Thus, with ROU, LC-Ve may paradoxically perpetuate pathophysiology through allostasis (Koob and Le Moal, 2001; McEwen, 2001).

While the stimulatory effect of fentanyl on LC activity did not substantially change with ROU, AVR and NVR mice diverged in their reliance on breathing outside of LC activity during OIRD. Whereas breathing outside of LC activity improved following ROU in AVR mice, it remained unchanged in NVR mice. Such an improvement in AVR mice consequently minimized the waxing and waning breathing pattern and lowered the gain of each LC-coupled event during OIRD. In NVR mice, a lack of improvement in Ve outside of LC activity preserves the gain of LC-coupled breathing observed when mice are naive to fentanyl exposure. These data suggest that LC-coupled breathing is a distinguishing feature of the AVR and NVR endotypes.

Respiratory endotype divergence aligns with the broader literature establishing that opioid responses vary across subjects. As documented previously (Rai et al., 2025), OIRD tolerance progressively emerges over the course of ROU in AVR mice, which represent the majority of subjects, and fails to develop in the NVR (Rai et al., 2025). While LC-Ve was not different between AVR and NVR mice on Day 1 of ROU, the gain of LC-Ve was reduced in the AVR mice after ROU. These data suggest that, as opposed to endotypes representing distinct groups a priori, the two endotypes mechanistically diverge as ROU progresses. Inbred mice in a shared environment progressively diverge into stable individual phenotypes attributable to stochastic and developmental influences (Freund et al., 2013). Such mechanisms may drive our observations, as the mice were from the same genetic background and underwent identical drug regimens. Opioid-experienced rats die at a greater frequency in response to a high dose of heroin in a context not previously paired with the drug rather than in the previously drug-paired context (Siegel et al., 1982), suggesting that opioid use leads to labile overdose tolerance with conditioned learning. Although Hou et al. (2023) described a similar phenomenon where pairing repeated morphine exposure with a salient context augments ventilatory tolerance to morphine, whether such tolerance remains in contexts with no prior drug association was not resolved. To address this, we incorporated salient cues into our protocol and tested whether the contextual pairing of fentanyl exposure impacted OIRD magnitude following ROU. Unexpectedly, we observed that OIRD magnitude was similar between the drug paired (FP) and unpaired (SP) environments. While LC-Ve differed between contexts in NVR mice, but not in AVR mice, the group × context interaction was not significant.. These results suggest that while context of drug use may influence the pattern of OIRD and the consequences of neuromodulation, OIRD tolerance, when acquired, is not affected by different salient contexts. Thus, further work is necessary for resolving the interaction between context and opioid overdose survival.

Our study and interpretations of the results warrant consideration of technical limitations and outcomes. First, our fiber photometry approach measures population Ca^2+^ dynamics and does not provide resolution from individual LC neurons. Thus, it remains possible that distinct ensembles in the LC are differentially recruited during OIRD. Second, although our endotype classification of AVR and NVR mice captured robust, reproducible divergence in breathing during OIRD following ROU, it remains unclear whether AVR and NVR are truly discrete categories or the extremes of a continuous distribution. For this reason, we have also provided the mean response without endotype stratification. However, given previously reported differences in ventilatory-metabolic coupling following ROU between AVR and NVR mice (Rai et al., 2025) and the potential for differences in ventilatory patterning during OIRD to perpetuate overdose status, we contend that stratification of AVR and NVR mice reflects genuine neurobiological behaviors and endotypes significant in understanding susceptibility to fentanyl overdose.

Independent of these considerations, our study identifies LC-coupled breathing as a physiological signature of variability in the OIRD response prior to and following ROU. This emerges even when animals have identical drug exposure histories and similar genetic backgrounds. Such heterogeneity carries direct relevance to overdose, as the strength and pattern of LC-driven breathing may indicate vulnerability to opioid-related death. Thus, our study also demonstrates that neuronal activity during OIRD continues to shape the pattern of breathing, opening the possibility that breathing during this state may serve as a physiological and behavioral biomarker of opioid susceptibility.

## Materials and methods

### Animals

All animal protocols followed National Institutes of Health guidelines for the appropriate care and use of animals and were approved by the Institutional Animal Care and Use Committee at The University of Chicago. Reporting follows the ARRIVE guidelines. For fiber photometry studies, we used a transgenic mouse line in which Cre-recombinase expression is driven by the norepinephrine (NE) transporter promoter and has high selectivity for NE cells in the locus coeruleus (LC) with low ectopic expression (slc6a2; mouse line Net-Cre; Wagatsuma et al., 2018). Adult mice of both sexes (>9 weeks, 22 - 41 g) on a Net-Cre background were used for simultaneous plethysmography and fiber photometry studies. Mice were bred at The University of Chicago animal facilities. Mice used in this study were group-housed with the same sex in AAALAC-approved facilities following a 12/12 h light/dark cycle and were given ad libitum access to food and water. All behavioral experiments occurred during their light cycle.

### Surgery

#### Stereotaxic surgical procedures

All viral injections were done with a stereotaxic surgical device (Kopf Instruments, Tujunga, CA, United States) to position injections and implants correctly, and sterile tools were used throughout to maintain a sterile surgical environment. Isoflurane (2 - 3% induction, 1 - 1.5% maintenance) was used to anesthetize the mice. Head hair was shaved using a trimmer, and the mice were head-fixed in the stereotaxic system. Sterile drapes were placed over a heating pad to maintain homeostatic body temperature, and ophthalmic lubricant was placed over the eyes. Betadine was applied on the scalp prior to any incision or injection. Bupivacaine (1 mg/kg, s.c.) was then injected into the scalp. Sterile tools were used to make an incision at the midline of the scalp. A micromotor drill attached to the stereotaxic apparatus was used to drill two small holes into opposite sides of the skull where two small screws were implanted. At the end of the surgery, mice were injected with meloxicam (5 mg/kg, s.c.) and saline (5 mL, s.c.) and returned to their cage. The cage was then placed on a heating tray until the mice recovered and regained normal mobility (about 1 - 3 days).

#### Viral injections and cannula implants for fiber photometry

A Hamilton syringe was attached to the stereotaxic apparatus for viral injections and slowly lowered to the locus coeruleus. A syringe pump was used to control the viral injection volume to 400 nL and flow rate to 100 nL/min. The needle was held in place for a total of 10 minutes to allow for sufficient viral infusion and was then slowly removed. To selectively monitor the activity of the noradrenergic cells in LC (LC^NE^), we injected a Cre-dependent genetically encoded Ca^2+^ indicator (GCaMP) virus (pGP-AAV-CAG-FLEX-jGCaMP8s-WPRE; Addgene 162380, Watertown, MA, United States) into the LC. Net-Cre transgenic mice were used for selective expression of GCaMP in LC^NE^ neurons. Fiber optic cannulas (MFC_400/430-0.66_6mm_MF1.25_FL; Doric Lenses, Quebec, Canada) were implanted using a cannula holder attached to the stereotaxic system. Dental cement was attached to the screws, skull, and cannula to create a head cap to hold and protect the fiber optic cannula. Experimenters waited at least 3 weeks for the virus to fully express before checking the fluorescent signal. LC coordinates relative to bregma for cannula placement were AP −5.4, ML 0.85, DV −3.75. Two DV viral injections were done dorsal and ventral to the coordinate at −3.9 and −3.6. DV cannula placement was at −3.75 (verified using Paxinos and Franklin, 2001).

### Behavioral protocols

#### Repeat opioid use protocol with context pairing

Both breathing with plethysmography and LC^NE^ activity with fiber photometry were measured during this protocol. The duration of the repeat opioid use (ROU) protocol was seven days and consisted of two phases. Phase One consisted of context pairing with either fentanyl or saline for five consecutive days. Adult mice expressing GCaMP in LC^NE^ were administered fentanyl (0.7 mg/kg, i.p.) daily in a distinctive environmental context (fentanyl-paired context). The same mice were also administered saline (0.2 - 0.3 mL, depending on mouse weight, i.p.) daily in another distinctive environmental context (saline-paired context). To make the environments distinctive, each had visual, tactile, audio, and olfactory cues associated with them. Because mice needed to remain in the plethysmography chamber to gather breathing data, the chamber became part of the environmental context. See-through chambers were placed into an opaque box to prevent mice from seeing experimenters. The box had visual cues on the walls and around the chamber, and audio that was played outside of the chamber. Scents for olfactory cues were mixed with the room air and flowed directly into the chamber; floor bottoms of the chamber were textured and acted as the tactile cue. Context one: green box with triangle taped on one side, four 3-D blue or purple shapes, floor bottom with small circular holes in it, white noise audio, and vanilla scent. Context two: white box with square and circle taped on two walls, four 3-D orange shapes, floor bottom made of mesh wiring, 80 BPM metronome audio, and almond scent. Which context was fentanyl-paired and which was saline-paired was counterbalanced across all of the subjects. Previous experiments demonstrated that there was no difference in the degree of respiratory depression depending on time of day of administration (e.g., morning or afternoon). To avoid any lingering effects of fentanyl carrying over into saline sessions, mice were injected with saline and placed in their saline-paired context each morning, and injected with fentanyl and placed in their fentanyl-paired context each afternoon. There was an average of 3.5 hours between the morning and afternoon sessions. Sessions consisted of a 15-minute baseline recording prior to any injection, and then a 15-minute recording after the injection of either fentanyl or saline. Phase Two consisted of a two-day protocol directly following Phase One. During Phase Two, there was only one session a day and mice only received fentanyl injections. Sessions again consisted of a 15-minute baseline recording prior to injection, and a 15-minute post-fentanyl injection recording. On one day of the protocol, mice again received fentanyl in the fentanyl-paired context; on the other day they received fentanyl in the saline-paired context. Which context came first during Phase Two was counterbalanced across the subjects.

### Immunohistochemistry and microscopy

Immunofluorescent staining was performed on all sections from a single series, at room temperature and with agitation, unless otherwise noted. Tissue was rinsed in 1X phosphate-buffered saline (PBS, pH 7.4) 3 times for 10 minutes each to remove cryoprotectant. Tissue was then washed in PBS supplemented with 0.5% Tween-20 (PBS-T) twice for 5 minutes each, and placed in a blocking buffer for 30 minutes (50 mM glycine, 0.1% (v/v) Triton X-100, 0.1% (w/v) bovine serum albumin, 0.05% (v/v) Tween-20, and 2% (v/v) normal donkey serum, in PBS). The primary antibodies rabbit anti-tyrosine hydroxylase (EMD Millipore AB152, Burlington, MA, United States; RRID:AB390204) and chicken anti-GFP (Invitrogen A10262, Waltham, MA, United States; RRID:AB2534023) were diluted 1:500 and 1:200, respectively, in antibody buffer (PBS supplemented with 0.1% Triton X-100, 0.05% Tween-20, 0.1% (v/v) hydrogen peroxide, and 10 mM glycine), which was applied to the tissue overnight at 4 °C. The next day, tissue was washed 3 times for 5 minutes each in 0.5% PBS-T, then incubated for 2 hours with secondary antibody solution, containing 1:1000 each donkey anti-chicken IgY (H+L) Alexa Fluor 488 (Invitrogen A78948) and donkey anti-rabbit IgG (H+L) Alexa Fluor Plus 647 (Invitrogen A32795) in 0.1% PBS-T. Afterwards, tissue was washed three times for 5 minutes each in PBS, mounted on glass slides, and coverslipped with ProLong Gold with DAPI (Invitrogen P36935).

Images were captured on a Zeiss Axio Imager.M2 upright epifluorescence microscope (Carl Zeiss, Oberkochen, Germany) using a 10x N-Achroplan 0.25 numerical aperture objective. All sections were imaged with identical collection parameters. Image files were first captured as z-stacks with a step size of 1 μm, and then flattened using the two-dimensional maximum intensity projection algorithm in ImageJ (NIH, Bethesda, MD, United States; RRID:SCR_003070) before quantification. Total cell counts within LC were made by hand for TH-positive or GFP-positive cells from the individual fluorescent channel images. Dual-labeled cells were quantified by using the Image Calculator function in ImageJ to create a co-localization mask based on fluorescent intensity overlap, and the output was similarly counted by hand. Solely single-labeled cell counts (e.g., TH-positive but GFP-negative) were calculated by subtracting the colocalized cell count from the total cell count within each section. All measurements were done first within single sections per subject, then averaged for all sections within a subject before further plotting and statistical analyses.

### Simultaneous whole-body plethysmography and fiber photometry

#### Whole-body plethysmography

Respiration was measured daily during the ROU protocol using unrestrained whole-body plethysmography in a custom-constructed chamber (500 mL) connected to a strain gauge pressure transducer. To conduct simultaneous fiber photometry recordings, the fiber optic patchcord was fed through a hole in the top of the chamber and attached to the mouse’s fiber optic cannula. An air-tight environment inside of the chamber was maintained by sealing the hole with modeling clay. Transducer signals were amplified via a MAX II amplifier (Buxco/Data Sciences International) and recorded using Synapse software (Tucker-Davis Technologies, Alachua, FL, United States; RRID:SCR006495). Room air (19-21% O₂, balanced N₂) continuously flowed through the system at 0.6 liters per minute. Chamber temperature (22-27 °C) was recorded each day of the ROU with context protocol prior to mouse placement in the plethysmography chamber. Barometric pressure was assessed across experimental sessions by using a website that provided the average barometric pressure (timeanddate.com) for Hyde Park, IL for each experimental day. Body weight and temperature (35.7-37.3 °C) were recorded for VT calculation, and mice were placed in the chamber and tethered to the fiber optic cable to record baseline breathing and LC^NE^ activity (15 minutes). At the end of the control period, calibrations for baseline VT were taken, the mouse was removed from the chamber and administered fentanyl (0.7 mg/kg, i.p.) or saline (0.2-0.3 mL, depending on animal weight) before being promptly returned to the chamber and re-tethered to the fiber optic cable to record drug-affected breathing and LC^NE^ activity (15 minutes). Post-injection calibration measurements for VT calculations were done prior to the removal of the mouse from the chamber at the end of the 15-minute fentanyl or saline recording. Once the animal was removed, body temperature measurements were again recorded (36.1-37.8 °C). Prior to the experimental protocol, mice were habituated to tethering, handling, i.p. injections (saline, 0.2-0.3 mL, depending on mouse weight), and a plethysmography chamber of a slightly different shape that would not be used in the protocol, for 5 consecutive days.

#### Fiber photometry

Fiber photometry was conducted using the TDT RZ10X system (Tucker-Davis Technologies, Alachua, FL, United States) with a lock-in amplifier for signal demodulation, in conjunction with a Doric 6-port fluorescent minicube (FMC6_IE(400-410)_E1(460-490)_F1(500-540)_E2(555-570)_F2(580-680)_S; Doric Lenses, Quebec, Canada). The minicube was equipped with 415 nm (isosbestic control) and 465 nm (GCaMP) excitation and 525 nm emission filters suitable for our fluorophores of interest. The 465 nm (GCaMP) channel was modulated at 330 Hz and the 415 nm (isosbestic control) channel was modulated at 210 Hz. The signal was sampled at 1017.3 Hz. Output power levels were matched between isosbestic and GCaMP signals and were kept consistent throughout the 7 days of the protocol. A Doric mono fiberoptic patchcord (MFP_400/430/1100-0.57_2m_FCM-MF1.25_LAF) was tethered to the fiber optic cannula on the animal’s head through a hole in the top of the plethysmography chamber. Mice were habituated to tethering during the 5-day habituation prior to any experimentation. TDT Synapse software was used to record and process fluorescent signals, track power levels, and simultaneously capture breathing signals. Photometry sessions lasted 30 minutes on average, including both baseline and fentanyl/saline sessions.

#### Post hoc analysis of simultaneous whole-body plethysmography and fiber photometry

Fiber photometry and breathing data collected through the TDT system were analyzed in MATLAB R2025b (MathWorks, Natick, MA, United States; RRID:SCR_001622) using the TDTbin2mat function. LC^NE^ activity and breathing were analyzed using a combination of MATLAB and custom scripts. The last 5 minutes of baseline were analyzed. The 15 minutes of fentanyl data were analyzed in 5-minute bins (minutes 0-5, 5-10, 10-15), to ensure proper fitting of 415 nm to 465 nm. Raw GCaMP data were corrected for motion artifact and baseline fluctuations by fitting the 415 nm isosbestic signal to the 465 nm GCaMP signal with the polyfit function. We then subtracted the 415 nm fit from the 465 nm signal and divided the difference by the 415 nm fit (ΔF/F = [F465 - Ffit415]/Ffit415). A Z-score of ΔF/F was used to compare across mice and across days. For both baseline and fentanyl/saline sessions, we used the mean and standard deviation from the baseline session of that day to calculate the Z-score.

The findpeaks function was used to locate LC fiber photometry events. The threshold for an event to be counted was set at a Z-score of 2 for all days, sessions, and animals. This is to ensure with 95% confidence that the events being counted were significantly different from the mean fluorescent signal. Various metrics were obtained about each LC event (amplitude, width, duration) and the breathing during that LC event (widths, amplitude, instantaneous frequency). The findpeaks function was used to identify breaths. Respiratory rate (RR, breaths·min^−1^) was calculated by multiplying the average instantaneous frequency by 60 seconds. Tidal volume (V_T_, μL·g^−1^) was calculated using the amplitude of breaths as described in Lim et al. (2014) and was normalized to weight. Baseline periods used baseline calibrations and body temperature recordings for V_T_ measurements, and post-injection (fentanyl or saline) periods used post-injection calibrations and body temperature recordings for V_T_ measurements. Minute ventilation (uL*g^-1^*min^-1^) was calculated as the product of tidal volume and respiratory rate. Breathing during the last five minutes of the 15-minute baseline period was analyzed and used as a baseline reference. Breathing was analyzed for the duration of the 15-minute post-injection period. To determine the relationship between LC activity and Ve, timestamps and durations of individual LC events were used to map back onto respiratory recordings and extract ventilation measurements during and 1 second before the start of the LC event.

#### OIRD ratio index

The response to ROU was calculated by the OIRD Ratio Index (ORI):

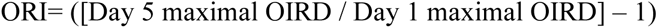

Where maximal OIRD was defined by the minimum Ve during fentanyl on Day 1 and Day 5, respectively.

The adaptive ventilatory response (AVR) following ROU was defined by a ORI > 0. The non-adaptive ventilatory response (NVR) following ROU was defined a ORI ≤ 0.

#### LC event-aligned analysis of ventilation

To examine the relationship between peak LC activity and ventilation, breath-by-breath Ve was aligned to the maximum of each detected GCaMP event from the LC. Analyses were performed using LC event timestamps and breath-level respiratory measurements generated from the post hoc pipeline described above. When two LC events occurred in close succession, they were merged if the interval between the end of the first event and the start of the next was less than or equal to 0.5 s. For merged events, the combined start and end times were retained, and the LC peak time was assigned to the event with the larger peak fluorescence amplitude. Breath-level Ve values were assigned to LC event windows using the absolute timing of each breath. Because Ve was measured breath-by-breath, each value was treated as constant over the corresponding breath interval. Outlier breaths were identified within each recording from the distribution of breath extrema and rejected if the maximum or minimum value exceeded three standard deviations from the recording-level distribution. For each LC event, Ve was normalized to the immediately preceding out-of-LC breathing period. The reference value was calculated as the time-weighted mean Ve during the 1 s window before LC onset, excluding rejected breaths and time overlapping with a preceding LC event. LC-associated Ve was expressed as the fractional change from this local reference and multiplied by 100 to indicate percent change from out-of-LC ventilation. LC-aligned matrices were generated by sampling normalized Ve from −0.6 to +0.6 s relative to the LC peak. LC events with incomplete coverage across the analyzed window were excluded from downstream analysis.

### Statistical analysis

Statistical analyses were performed using GraphPad Prism 10 (GraphPad Software, Boston, MA, United States; RRID:SCR002798), with data expressed as means and significance set to p < 0.05, or custom scripts written in Python 3.10.17 (RRID:SCR008394) using statsmodels 0.14.4 (RRID:SCR_016074). Two-condition repeated measures comparisons were evaluated by paired t-test or Wilcoxon test. Data that were not normal were analyzed using non-parametric method (Wilcoxon matched pairs). Unpaired t-test was used to compare data not temporally related. Two-factor designs were assessed using RM two-way ANOVA or Mixed Effects analysis with Tukey’s multiple comparison and Geisser-Greenhouse correction or Fisher’s LSD. Time-resolved statistical comparisons were performed using linear mixed-effects models at each LC-relative time point. For D1 versus D5 and SP versus FP comparisons, day or context was included as a fixed effect, and animal identity was included as a random effect. To test whether LC-associated ventilatory responses differed between AVR and NVR mice, response group and the day-by-group or context-by-group interaction were included as fixed effects, with animal identity included as a random effect. Statistical significance across time was assessed using cluster-based permutation testing. Clusters were defined as contiguous time points with absolute model t-values greater than 2, and condition labels were permuted within each animal for 1000 permutations to generate the null distribution of maximum cluster statistics. Clusters with permutation-corrected p < 0.05 were considered significant. Outliers were identified using ROUT test (Q = 1%) and subsequently excluded. Normality was tested using Shapiro-Wilk or Kolmogorov-Smirnov tests where appropriate for sample size.

## AI use disclaimer

Generative AI tools were used during the preparation of this manuscript for (i) language editing and revision of author-written text, (ii) checking reference accuracy and formatting, and (iii) assisting in the development of custom analysis code (MATLAB and Python) used to process and analyse plethysmography and fiber photometry data. AI-generated code was reviewed, tested, and validated by the authors. We take full responsibility for the analytical pipelines and the results derived from them. AI tools were not used to collect data, interpret results, or generate figures. All AI-assisted text was reviewed and edited by the authors.

## Materials availability

All biological reagents and custom analysis code described in this article are available from the corresponding author upon reasonable request from academic researchers, without restriction on use.

## Acknowledgements

The authors thank members of the Institute for Integrative Physiology and The Neuroscience Institute at The University of Chicago for helpful discussion.

## Additional information

### Competing interests

The authors declare that no competing interests exist.

### Funding

5T32DA043469 (CCS); The University of Chicago, Safadi Pilot Grant (AJG); R01HL163965 (AJG); R01DA057767 (AJG); R01DA061412 (AJG); R01HL169679 (AJG)

### Author contributions

CCS and AJG conceived and designed experiments; CCS, MW, JVG, SS performed experiments; JSP, AY, GEF conducted analyses; JSP and AJG drafted the manuscript; all authors revised the manuscript. AJG provided reagents and support.

### Ethics

All animal protocols followed National Institutes of Health guidelines for the appropriate care and use of animals and were approved by the Institutional Animal Care and Use Committee at The University of Chicago.

### Data availability

All source data and analysis codes can be made available upon reasonable request.

---

**Supplement Figure 1.**
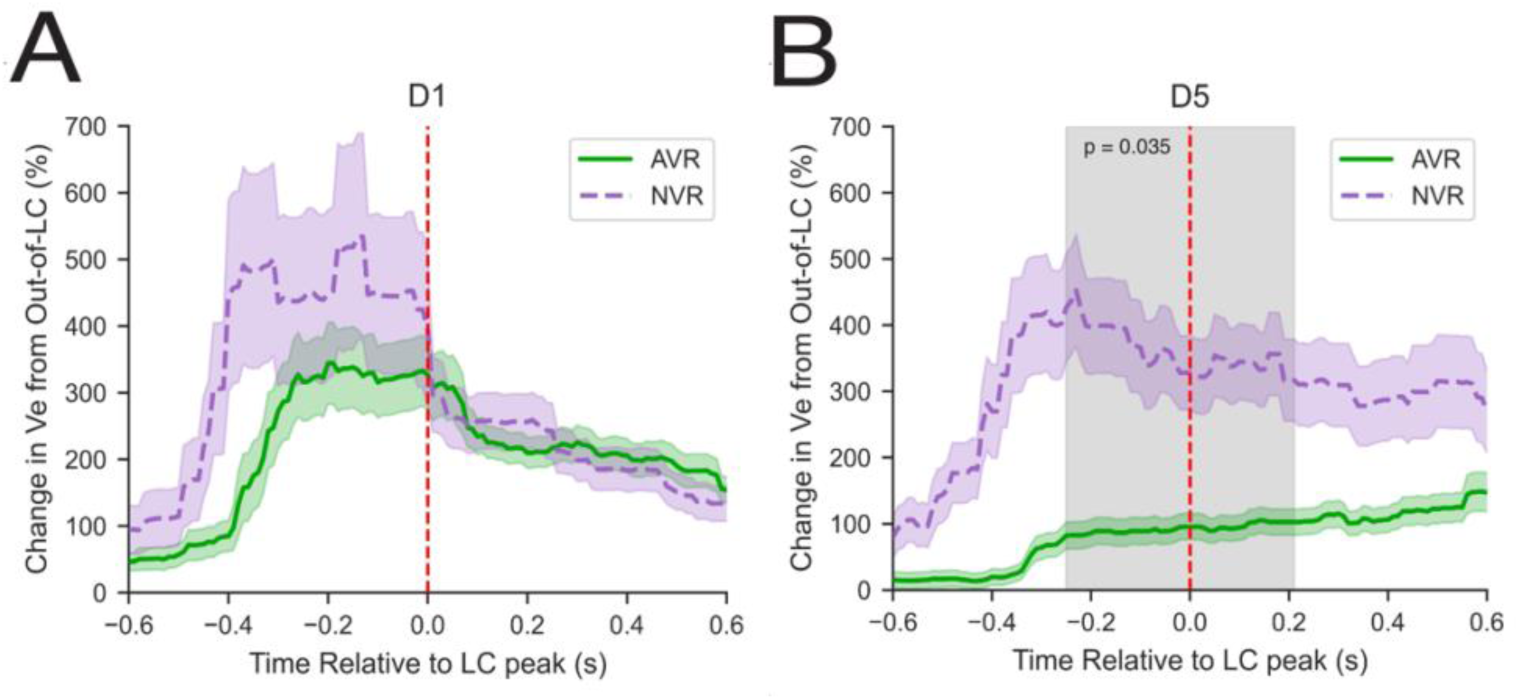
Respiratory endotypes show divergent LC-Ve during OIRD following ROU. (A) Time course of Ve relative to peak LC activity during OIRD from AVR (n = 6) and NVR (n = 3) groups on Day 1 shows no difference between respiratory endotypes prior to ROU. (B) Time course of Ve relative to peak LC activity during OIRD on Day 5 from AVR and NVR groups (p = 0.035). (A, B) Linear mixed-effects model, significance determined by cluster-based permutation test. Shaded area indicates significant difference between groups.

**Supplement Figure 2.**
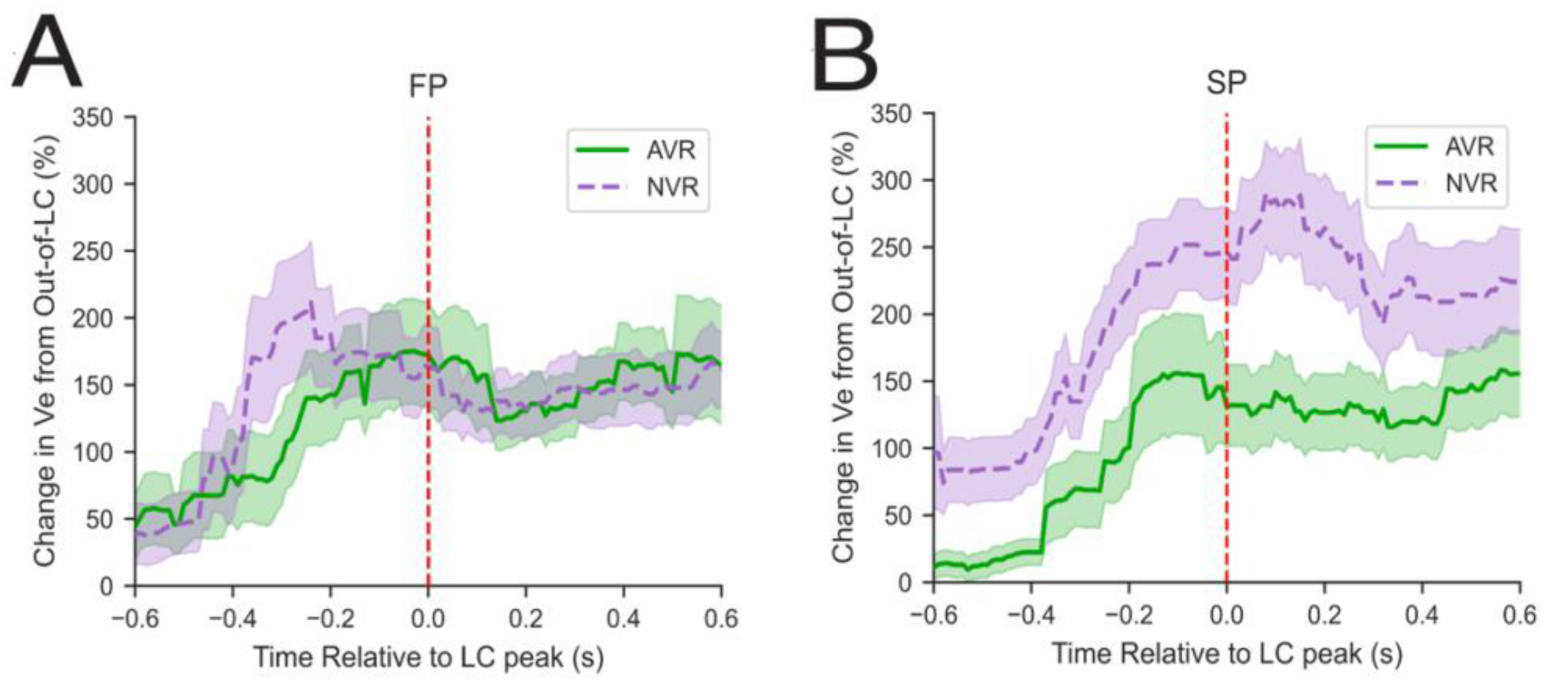
LC-Ve throughout OIRD from respiratory endotypes in SP and FP contexts. (A) Time course of Ve relative to peak LC activity during OIRD from AVR (n = 6) and NVR (n = 3) groups in the FP context shows no difference between respiratory endotypes. (B) Time course of Ve relative to peak LC activity during OIRD in the SP context from AVR and NVR groups. (A, B) Linear mixed-effects model, significance determined by cluster-based permutation test. Shaded area indicates significant difference between groups.

## Notes

### Competing Interest Statement

The authors have declared no competing interest.

